# Demographic inference provides insights into the extirpation and ecological dominance of eusocial snapping shrimps

**DOI:** 10.1101/2020.09.07.283994

**Authors:** Solomon T. C. Chak, Stephen E. Harris, Kristin M. Hultgren, J. Emmett Duffy, Dustin R. Rubenstein

**Author notes:** Corresponding author: Solomon T. C. Chak, Department of Biological Sciences, SUNY College at Old Westbury, Old Westbury, NY 11568, USA. Both authors contributed equally to this manuscript (Solomon T. C. Chak, Stephen E. Harris).

## Abstract

Eusocial animals often achieve ecological dominance in the ecosystems where they occur, a process that may be linked to their demography. That is, reproductive division of labor and high reproductive skew in eusocial species is predicted to result in more stable effective population sizes that may make groups more competitive, but also lower effective population sizes that may make groups more susceptible to inbreeding and extinction. We examined the relationship between demography and social organization in one of the few animal lineages where eusociality has evolved recently and repeatedly among close relatives, the *Synalpheus* snapping shrimps. Although eusocial species often dominate the reefs where they occur by outcompeting their non-eusocial relatives for access to sponge hosts, many eusocial species have recently become extirpated across the Caribbean. Coalescent-based historical demographic inference in 12 species found that across nearly 100,000 generations, eusocial species tended to have lower but more stable effective population sizes through time. Our results are consistent with the idea that stable population sizes may enable eusocial shrimps to be more competitively dominant, but they also suggest that recent population declines are likely caused by eusocial shrimps’ heightened sensitivity to anthropogenically-driven environmental changes as a result of their low effective population sizes and localized dispersal, rather than to natural cycles of inbreeding and extinction. Thus, although the unique life histories and demography of eusocial shrimps has likely contributed to their persistence and ecological dominance over evolutionary timescales, these social traits may also make them vulnerable to contemporary environmental change.

---

Eusocial species are often the most abundant members of their community (1, 2). Their ability to outcompete conspecifics, expand their niches, and assert their ecological dominance over non-eusocial species—an idea referred to as the ‘social conquest hypothesis’—may be due in part to their ability to cooperate and form complex societies (2, 3). Wilson (4) proposed several population-level qualities of life history and demography that contribute to the ecological success of eusocial ants and their persistence over evolutionary time. Since all of the more than 13,000 species of ants are eusocial (5), testing these ideas empirically has been challenging. Moreover, whether these demographic characteristics are shared in other eusocial organisms that exhibit various forms of social complexity and ecological success remains unclear. Although a few studies have examined empirically and comparatively whether social populations and eusocial species have been able to expand their ecological niches (3, 6, 7) and geographic ranges (8) relative to non-social ones, the specific life history and demographic characteristics that allow eusocial species to maintain high population densities that can be sustained for long periods of time remain largely unexplored.

Social conquest and ecological dominance may be facilitated or hindered by several life history characteristics specific to eusocial species that are known to influence demography and genetic diversity. First, in eusocial species, reproductive division of labor and high reproductive skew may lead to a reduction in effective population size (N_e_) (9, 10) because the breeding population (reflected by N_e_) is constrained to mainly a single or few reproductive individuals (i.e. queens). In contrast, a much larger portion of the population contributes to breeding in non-eusocial species. Second, reproductives in eusocial species may be buffered from environmental fluctuation by the workers that cooperatively attend them, resulting in more stable N_e_ over time (11, 12). Similarly, the ability to cooperatively defend and maintain a stable domicile (4) may further buffer the breeding individuals—and indeed the entire group—from environmental fluctuation, similarly resulting in more stable N_e_ over time. Together, the demographic consequences of these life history characteristics are likely to be lower N_e_, but greater population stability over time in eusocial species than non-eusocial species (4). Yet, at the same time, these or other demographic correlates of eusociality can also render some eusocial lineages unstable, resulting in inbreeding, local extinction, and even evolutionary dead-ends. For example, lineages containing social spider species have undergone repeated origins and extinctions, likely due to inbreeding, reduced N_e_, and the depletion of genome-wide genetic diversity (13–15). Indeed, instances of inbreeding that generate low genetic diversity and the potential for colony collapse (16)—often in the face of environmental fluctuation—are surprisingly common in other primitively eusocial species like wood-dwelling termites, thrips, spider mites, and naked mole-rats (reviewed in 15). Ultimately, conflicting empirical evidence from insects lead to contrasting predictions about the role of historical demography in social evolution: reproductive division of labor and protection of female reproductives by workers predicts a historical trend of population stability in eusocial species, whereas low N_e_, the potential for inbreeding, and reduced genome-wide genetic diversity predicts long-term population instability for eusocial species.

Testing these contrasting predictions and bridging the gap between these alternative scenarios of the demographic stability of eusociality can be done using coalescent-based methods that infer historical demography. Historical expansions or contractions in population size, as well as population structure and gene flow, leave a measurable signal on the genetic diversity of contemporary populations, which allows for the quantification of the demographic process based on the site frequency spectrum (SFS), a measure of allele frequencies across the genome that can be generated from large-scale sequencing. The effects of demographic processes on the SFS are well studied and have been tested across a range of taxa (17–19). Demographic inference is commonly made using a composite likelihood framework that fits the observed SFS to the expected SFS computed under given demographic histories (17, 20–24). This approach has been used successfully in species ranging from mice (25) to humans (26). Recent studies have also begun modelling comparative demographic processes across multiple taxa simultaneously (20, 27–30), including eusocial bees (31) and social spiders (14). These and other studies examine relative patterns of N_e_ change through time, rather than simply comparing absolute estimates at specific time points (32–41), an approach that would enable us to test alternative demographic scenarios in eusocial lineages. Moreover, the broad use of coalescent-based methods has shown its robustness as a population genetic tool and can accurately select models and estimate parameters using as few as 3-15 individuals and 10,000 – 50,000 single nucleotide polymorphism (SNP) loci (42, 43). Therefore, it is now feasible to use a multi-taxa comparative demographic framework to test long-standing hypotheses underlying how population dynamics influence the ecological consequences of eusociality.

Here, we use a group of socially diverse, sponge-dwelling snapping shrimps in the genus *Synalpheus* to look at the demographic consequences of eusociality and to address whether historical N_e_ supports a model of demographic stability or instability in eusocial species. *Synalpheus* shrimps in the West Atlantic *gambarelloides* group represent a relatively young lineage that radiated between approximately 5 and 7 Mya (44), yet eusociality has evolved independently at least four times in this group (45) and likely represents an early stage of eusocial evolution because workers are not sterile (46, 47). Species in this clade exhibit a variety of forms of social organization, ranging from pair-living, to communal breeding (multiple mating pairs in the same sponge), to eusociality (one or a few queens and a larger number of non-sterile workers of both sexes) (45–48). Most importantly, there are reports of both ecological dominance and demographic instability in eusocial *Synalpheus* species. The diversity of *Synalpheus* species in Carrie Bow Cay, Belize has been well documented since 1990 and represents the only longitudinal dataset on snapping shrimps currently available. At this reef, the four eusocial shrimp species contributed to >65% of the quantitative samples and eusocial species were 17 times as abundant as non-eusocial species (49). Eusocial species also occupied more sponges and had a wider host breadth than their non-eusocial relatives (7). Thus, for more than two decades, eusocial *Synalpheus* outcompeted their non-eusocial relatives and remained the ecologically dominant shrimp species on the reef. However, since 2012, three of the four eusocial species at Carrie Bow Cay have become locally extinct. In addition, two eusocial species in Jamaica have shown population declines associated with an increase of queenless colonies (50), and a eusocial species in Panama has also declined (45). In these same reefs in Jamaica and Panama, pair-living and communal breeding *Synalpheus* species appear to at least be maintaining healthy populations, if not increasing in abundance. Therefore, despite evidence of sustained ecological dominance, eusocial *Synalpheus* shrimps have seen recent population declines over much of their range. These recent population declines could be a snapshot of cyclic population bottlenecks that reflect the long-term demographic instability of eusocial species. However, since the colony organization of eusocial species (i.e., reproductive division of labor and high reproductive skew) (4, 9) also predicts that they could have low but stable population sizes, if eusocial species show a historical trend of population stability, then these recent population declines are more likely to be explained by contemporary environmental change (e.g. increased anthropogenic environmental disturbance).

To explore the relationship between demography and eusociality, we generated genome-wide SNP data using double digest restriction-site associated DNA sequencing (ddRADseq) and performed demographic inference to compare long-term demographic patterns in four eusocial and eight non-eusocial shrimp species (including five communal breeding and three pair-forming species). We inferred historical N_e_ by finding which pattern of population size changes over time generated an expected SFS that best matches our observed SFS. Using inferred historical N_e_, we then calculated average N_e_ over multiple timepoints, assuming the same mutation and recombination rates across species (51), and three population stability metrics: (i) the coefficient of variation of N_e_ across times (CV) to quantify the variation of N_e_ that is independent of mean N_e_, where a smaller value indicates greater population stability; (ii) the ratio of minimum N_e_ to mean N_e_ (min/mean N_e_), where a larger value indicates a less severe population bottleneck, and thus greater population stability; and (iii) the number of times that two consecutive N_e_ estimates had less than one order of magnitude difference (no. of <1-order change), where a larger value indicates less drastic changes in N_e_, and thus greater population stability. Ultimately, linking demography to social evolution through population genetic approaches (52) not only has the potential to uncover the demographic factors that may have enabled some eusocial species to come to dominate the ecosystems where they occur, but it may also help explain why some eusocial species or lineages go extinct over both evolutionary and ecological timescales.

## Results

### Genetic diversity and kinship

For each of the 12 *Synalpheus* species, we sampled 7-8 breeding females (i.e., queens) from different sponges (i.e., colonies) from the same reef whenever possible. Using ddRADseq, we generated >312 million paired-end reads with a median of 2.7 million reads per sample. De novo mapping using *Stacks* generated 19,788 – 52,898 SNPs per species (median = 24,307, Table S1 and S2; SRA accession: SAMN14351547 - SAMN14351641). Mean individual observed heterozygosity (H_obs_) based on all sites or just variant sites did not differ among species exhibiting the three forms of social organization (pMCMC > 0.19, Fig. S1, Table S3, Table S4): the percentage of heterozygous sites across all sites ranged from 0.19 to 0.23% across species, which is similar to that observed in non-social spiders (14) and other *non-Synalpheus* shrimps (53). Moreover, there were low levels of inbreeding in all species: the inbreeding coefficients (F) were positive and ranged from 0.12 to 0.54 across species (Fig. S1, Table S3). In contrast to the prediction that eusocial species may be more inbred than non-eusocial species (14), eusocial *Synalpheus* species were significantly more outbred than communal breeding species (pMCMC = 0.032), but only marginally more outbred than non-eusocial (pair-living and communal breeding combined) species (pMCMC = 0.062) (Table S4). Finally, mean average kinship coefficients were negative for all species, meaning that breeding females across sponges were unrelated to each other (Fig. S1, Table S3). However, kinship coefficients were significantly higher in eusocial species than in pair-forming species and non-eusocial species (pMCMC = 0.0003 and 0.02, respectively; Table S4). The relatively higher degree of genetic relatedness among queens over small spatial scales in eusocial versus non-eusocial species is consistent with their different modes of larval dispersal: all eusocial species have crawling larvae that remain in the natal sponge, whereas communal breeding and pair-living species have swimming larvae that disperse from the natal sponge via the water column (7, 45, 54, 55).

### Demographic inference and stability

Rather than simply comparing N_e_ values at specific time points, we examined relative changes in N_e_ between species to identify broad demographic patterns within groups of species exhibiting different forms of social organization. This approach avoided interpretation of the timing of events and absolute N_e_ values, both of which can be confounded by potential variations in mutation and recombination rates across species (51). Knowledge of mutation rates and generation times are not well known in *Synalpheus* shrimps and most other social animals (56), and thus, such an analysis could generate uncertainty in absolute N_e_ estimates. However, in lieu of exact point inferences, the main features of the model (i.e., relative changes, overall size changes, directionality, and broad patterns in demographic history) can be accomplished with confidence through bootstrapping simulated data under the best-model demographic scenario (57). Thus, due to the uncertainty in determining absolute N_e_ values, reporting on the broad demographic trends and relative changes in N_e_ is the accepted approach when using coalescent-based modeling to estimate demography from the site frequency spectrum (35–40, 58–61).

After removing related individuals (n = 5 of 96) and drawing one random SNP per locus, we used 11,890 – 26,148 SNPs per species (median = 18,175, Table S1) for demographic inference analysis using *MOMI2*, an efficient and scalable demographic inference software that can handle complex datasets (23, 24). For each species, we ran multiple analyses that varied in the number of times N_e_ was inferred (T = 4, 6, or 8 time points) and over different timescales (G = 60, 80, or 100 thousand generations). To examine N_e_ across species with the same underlying parameters (T and G), we compared the delta AIC (62) across analyses and found that a model that estimated N_e_ four times across 100,000 generations had the best support across species (Table S4). Estimated values of N_e_ from this model were generally within the 95% confidence interval of the bootstrap estimates (see Supporting Information: Fig. S2), indicating that the patterns were consistent across 300 different subsets of SNPs. Assuming several generations per year, a time scale of roughly 100,000 generations is likely to span most of the Holocene.

In general, N_e_ values were higher and more variable through time in pair-living and communal breeding species, whereas they were lower but more stable through time in eusocial species (Fig. 2). There appeared to be a reduction of N_e_ at 51-75 thousand generations ago for three out of four eusocial species. Similarly, three communal species also showed a reduction of N_e_ at 26-50 thousand generations ago. These shared trends indicate that these species may respond to the same demographic event and gave additional confidence to our estimates. However, since our primary aim was to examine the relative trends of historic N_e_ among species exhibiting different forms of social organization, in addition to estimating the average population size over multiple timepoints, we also calculated three population stability metrics using inferred historical N_e_ from the best fit model with the highest log-likelihood. We then used phylogenetic mixed models implemented in *MCMCglmm* (63) to compare these demographic metrics using both a continuous measure of eusociality (eusociality index) and across discrete social categories (64). Importantly, the eusociality index captures the reproductive skew within a group (7, 65, 66), making no *a priori* assumption of social phenotype. Our results confirmed the above patterns statistically in 3 of 4 metrics of population size and stability: as eusociality index increased (hence stronger reproductive skew), mean N_e_ decreased, CV decreased, and min/mean N_e_ increased, though no. <1-order change did not change (pMCMC = 0.018, 0.031, 0.025, and 0.63, respectively, Fig. 3; see Supporting Information Table S5). We observed qualitatively similar results using discrete social categories, such that eusocial species had significantly lower mean N_e_ than non-eusocial species (pMCMC = 0.021), as well as marginally lower CV, higher min/mean N_e_, and higher no. of <1-order difference in N_e_ values than non-eusocial species (pMCMC = 0.055, 0.058, and 0.076, respectively, Fig. 4; see Supporting Information, Table S5). Although the limited number of species in each social category likely contributed to the marginal levels of statistical significance, the results are consistent with those from the analysis using the eusociality index. Finally, analyses using the median demographic metrics across bootstrap estimates showed similar trends, but lacked statistical significance (see Supporting Information, Fig. S3), likely because smaller subsets of SNPs were used in these analyses. Thus, eusocial shrimp species have lower but more stable N_e_ through time than their non-eusocial relatives.

**Fig. 1.**
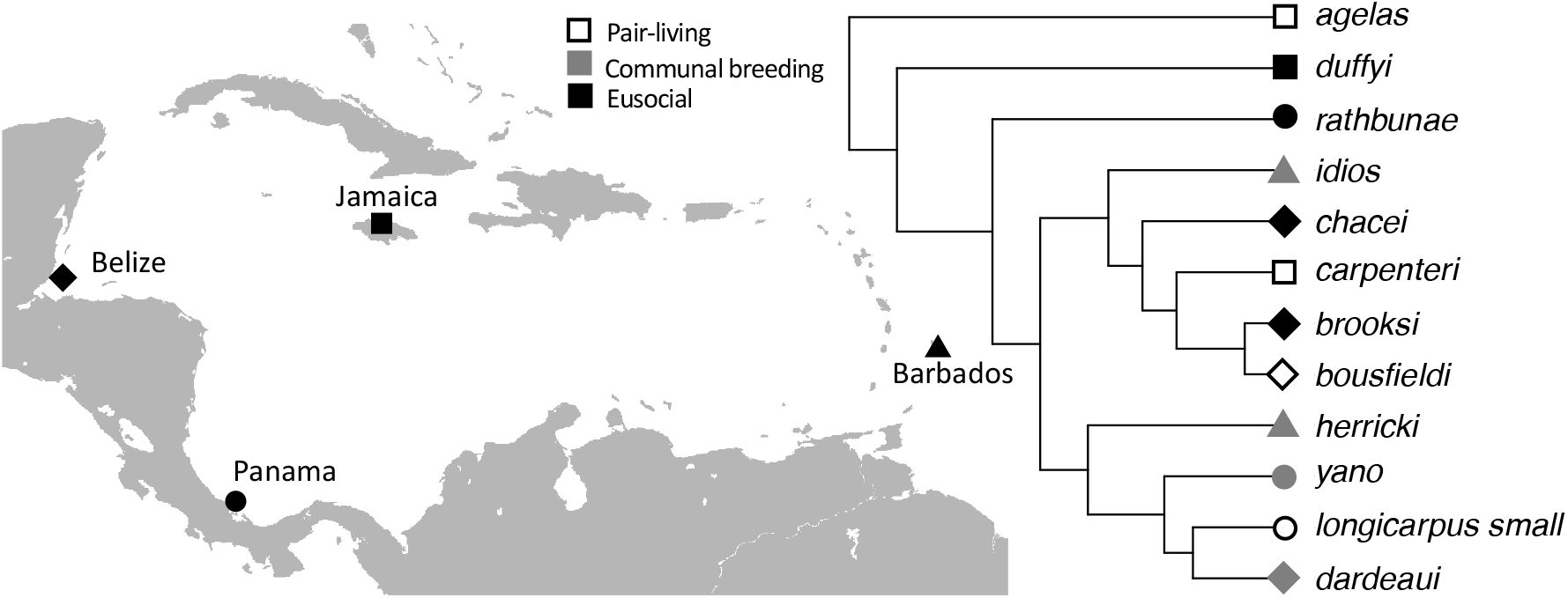
Collection sites and evolutionary relationships among 12 sponge-dwelling snapping shrimps in the genus *Synalpheus* used for comparative demographic inference. Each shape corresponds to a different sampling locality. The color of each shape corresponds to the form of social organization (white: pair-living, grey: communal breeding, and black: eusocial). The map is modified from Wikimedia Commons under CC BY-SA 3.0.

**Fig. 2.**
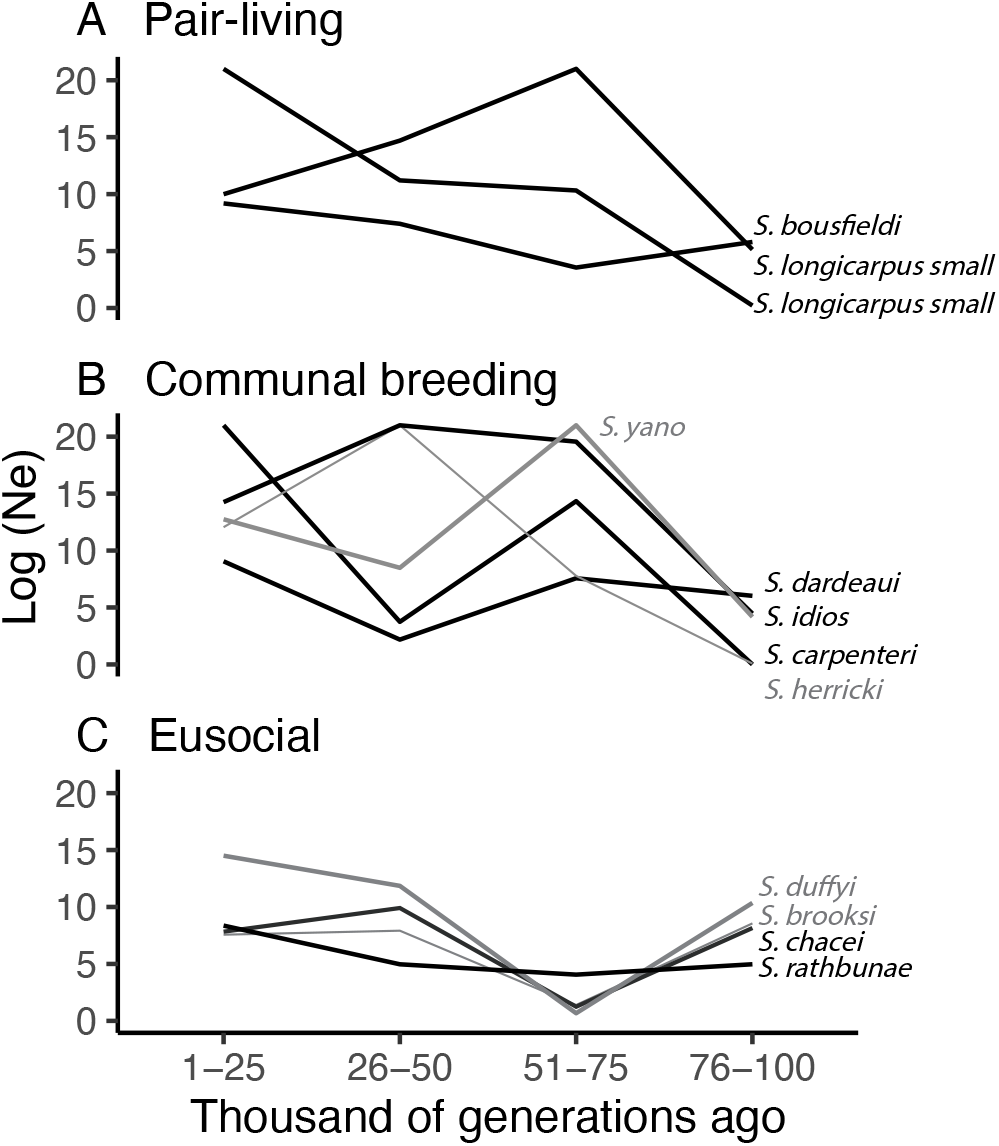
Best model estimates of N_e_ across 100,000 generations in 12 *Synalpheus* species exhibiting different forms of social organization (A: pair-living, B: communal breeding, C: eusocial). Each line represents N_e_ for a species, with species label on the right. Some lines were shown as grey or thin grey lines for clarity. Bootstrap estimates are shown in Fig S2.

**Fig. 3.**
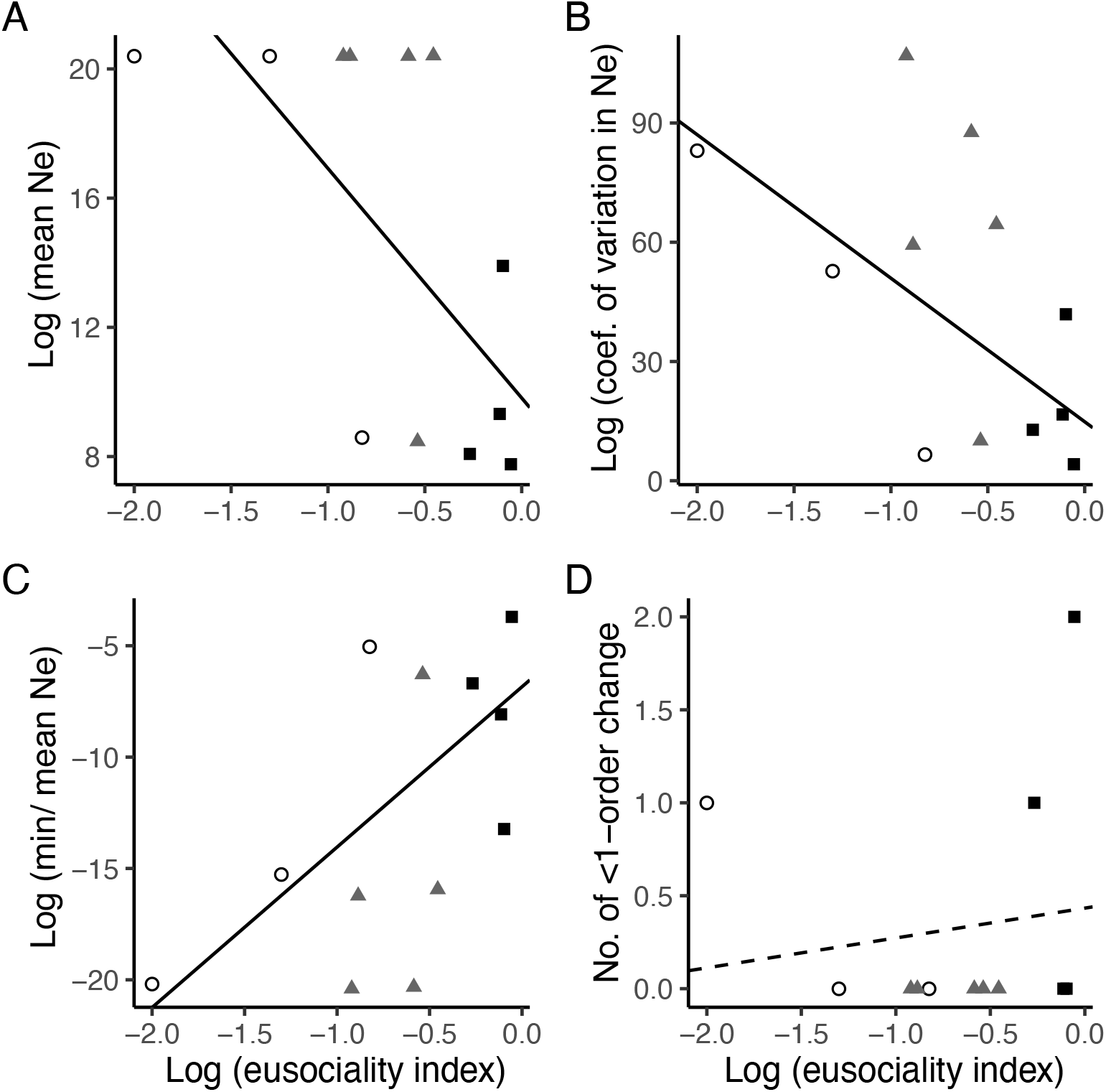
Relationships between the eusociality index and metrics of population size and stability in 12 *Synalpheus* species: (A) mean N_e_ across time, (B) coefficient of variation in N_e_, (C) min/mean N_e_ and (D) No. of <1-order change in N_e_. Symbols represent raw values: open circles = pair-forming species, grey triangles = communal breeding species, dark squares = eusocial species. Solid and dashed lines represent significant and non-significant regression slopes predicted using Bayesian phylogenetic mixed models, respectively.

**Fig. 4.**
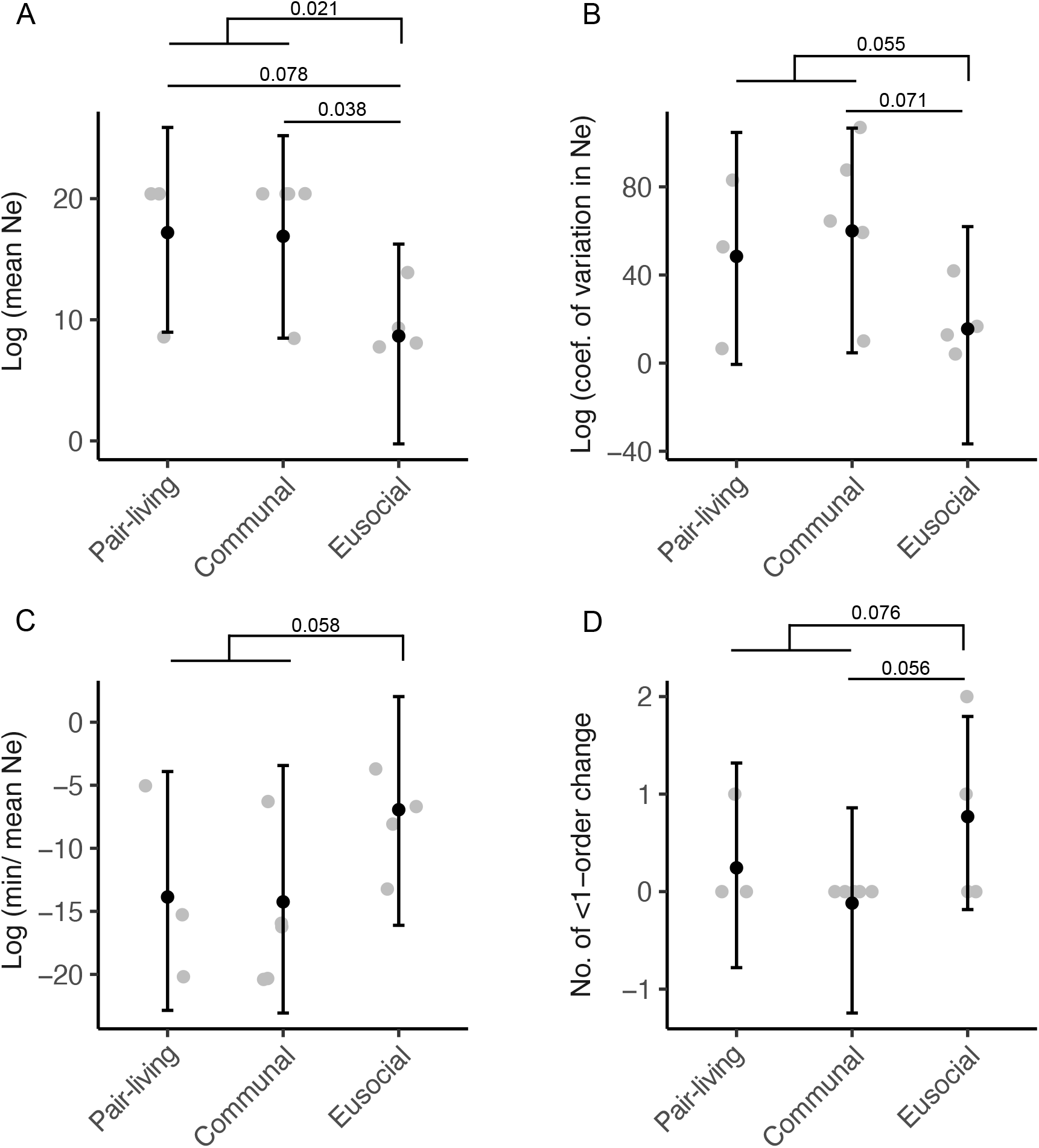
Metrics of population size and stability across 12 *Synalpheus* shrimps exhibiting different forms of social organization. Eusocial species have lower mean N_e_ across time (A), but more stable values of N_e_ across generation time as indicated by lower CV (B), higher min/mean N_e_ (C), and higher no. <1-order change (D). Grey dots are raw values, black dots are posterior mean predicted using Bayesian phylogenetic mixed model, bars are the 95% posterior distributions, and pMCMC values are twice the probability that the posterior distribution of the difference is above or below zero.

## Discussion

Cooperation between individuals of the same species leading to the formation of complex societies is thought to have enabled eusocial animals to become ecologically dominant in the areas where they occur, sometimes comprising more than half of the biomass in a given ecosystem (2). The peculiar demographic consequences of eusociality have been suggested to both help and hinder their ecological dominance, although empirical tests of these ideas have largely been lacking. Here, we examined the relationship between demography and eusociality in a socially diverse clade of *Synalpheus* snapping shrimps using historical demographic inference to determine whether eusocial species are more or less demographically stable than their non-eusocial relatives (4). We found that eusocial *Synalpheus* snapping shrimps, across four independently-evolved eusocial lineages, consistently had lower but more stable N_e_ through time than non-eusocial species. This long-term stability of N_e_ in eusocial shrimp species supports the idea that reproductive division of labor may enable groups to buffer the breeding individuals (i.e., N_e_) against environmental fluctuation or that cooperation may help eusocial species to better defend and maintain their domiciles. Furthermore, Wilson (4) argued that a critical criterion for ecological success was population density that could be sustained for long periods of time. Our results are consistent with this idea, adding to field observations that, for more than two decades, eusocial *Synalpheus* outcompeted their non-eusocial relatives to become the ecologically dominant shrimp species on the reef (7, 49). Although species with low mean N_e_ may be more susceptible to extinction due to reductions in genetic diversity and high genetic load (67), this does not seem to be the case in eusocial shrimp because eusocial species exhibited less evidence of inbreeding than non-eusocial species. Moreover, we found no evidence of population bottlenecks, cyclical patterns of expansion, or contractions of N_e_ in eusocial shrimp species, as found in other primitively eusocial animals (15). Instead, eusocial shrimp species appear to be more demographically stable than non-eusocial species, a conclusion that remains robust whether we consider social organization as a categorial or continuous variable. Our results agree with recent ecological models showing primitive eusociality to be generally more successful than solitary nesting in terms of reproduction and extinction probabilities (68). The evolutionary dead-ends observed in social spiders and other primitively eusocial animals may be explained by the advantages of solitary nesting compared to eusociality under very restricted assumptions in simulation studies (68, 69).

One of the primary motivations for this study was to quantify the demographic consequences of eusociality in snapping shrimps in order to understand the mechanism underlying the ecological dominance of eusocial species on Caribbean reefs. Yet, this study also has more practical implications since eusocial *Synalpheus* species in Belize, Panama, and Jamaica have either gone locally extinct or declined significantly in abundance in recent years, while non-eusocial species have remained stable or increased in abundance over the same timeframe (50). Our findings that eusocial *Synalpheus* species had stable effective population sizes over evolutionary time, and the absence of any evidence of population bottlenecks or cyclical patterns of expansion or contractions of N_e_, together suggest that these recent population declines are unprecedented in the last 100,000 generations, spanning most of the Holocene. We propose instead that contemporary population declines may be due to recent environmental changes across the Caribbean, which likely impact eusocial species more strongly because of their lower average N_e_ and limited ability to disperse. For several decades, direct anthropogenic development and environmental disturbances have negatively impacted coastal and marine ecosystems across the Caribbean (70). A combination of warming sea surface temperatures (71), rising sea level (72), and increased hurricane activity have been affecting the Caribbean, all of which negatively impact benthic communities (73, 74). The global increase in CO_2_ has also caused increased acidification, especially in the Caribbean region over the past 25 years, negatively affecting a wide range of biological systems (75). Together, these climate events have had devastating impacts on local ecosystems, including on sponge communities (e.g., 76). Since *Synalpheus* are obligate sponge dwellers, and many are strict host specialists (6), these symbionts may be particularly sensitive to environmental change. Within sponges that host *Synalpheus* shrimps, extreme environmental perturbations are likely driving the global increase in sponge diseases, which has especially affected Mediterranean and Caribbean sponges (77). Increasingly disturbed by anthropogenic pressures, the survival of local snapping shrimp populations is likely dependent on their ability to quickly and efficiently recolonize host sponges. However, the life history of eusocial shrimp species may hamper their ability to recolonize host sponge after a local population crash: eusocial shrimps tend to be host specialists (6) and all eusocial species have crawling larvae that remain in the natal sponge, in contrast to communal breeding and pair-living species that have swimming larvae that can disperse from the natal sponge in the water column (7, 54, 55). Our genetic diversity results support this life history difference: eusocial shrimps have the highest levels of relatedness (Fig. S1) across females from different sponges within a local population (collection site), a pattern seen previously using allozymes (78). Therefore, eusocial species appear to have lower migration and colonization rates than non-eusocial species, hence potentially less gene flow between populations, as shown in an early population genetics study (78). Thus, although eusocial *Synalpheus* species have been ecologically dominant in parts of the Caribbean for at least a few decades (7, 49) and demographically stable for thousands of years, in the face of increased anthropogenic environmental disturbance, eusocial species may be more susceptible to population collapse and extinction than their non-eusocial relatives.

In conclusion, our analyses are consistent with the idea that eusocial animals tend to be more demographically stable than their non-eusocial relatives; we found a signature of lower and more stable N_e_ in eusocial shrimps compared to non-eusocial species. In contrast, we found no evidence of cyclic population contractions or extinctions in eusocial species that would indicate population instability. Our results suggest that for at least 100,000 generations spanning thousands of years, eusocial shrimp populations in the Caribbean appear to have been more stable than those of non-eusocial species, suggesting that the recent extirpation of ecologically dominant eusocial species may be driven by anthropogenic changes in combination with their lower N_e_ and limited dispersal ability. Therefore, the long-term demographic stability in eusocial snapping shrimp is likely one of the key factors in promoting their ecological dominance on tropical reefs, an idea that has been proposed but not tested for eusocial insects that have come to dominate much of the terrestrial world. Yet, the demographic and life history characteristics of eusocial shrimps also render them vulnerable to recent global change. Ultimately, studying changes in demography through time will not only help understand the long-term population dynamics of social organisms, it may also provide critical insights into how social species will cope in the face of increasing climate change and habitat loss.

## Materials and Methods

### Sampling and study species

We sampled 12 species of *Synalpheus* snapping shrimps from four sites in the Caribbean (Belize, Barbados, Jamaica, and Panama) between 2004 and 2012 (Fig. 1). A detailed collection protocol has been reported previously (49). For each species, we sampled 7-8 females, enough for confident demographic inference (42, 43), with eggs or visible ovarian development to ensure that we were sampling breeding individuals. We sampled females from different sponges whenever possible. For eusocial species, each sponge represented a colony. Thus, sampling from different sponges ensured that we had a more complete representation of the population at each site. All individuals were collected within a 5-year sampling period (Supporting Information: Table S1).

Our sampled species included three pair-forming, five communal breeding, and four eusocial species. The four eusocial species included one species from each of the lineages that independently evolved eusociality in the genus *Synalpheus*. Two eusocial species (*S. brooksi* and *S. chaceĩ*) were from Belize where, before the recent population decline, they dominated their habitat over non-eusocial species based on abundance, sponge occupancy, and host ranges (7). Local population declines were observed from field surveys for *S. chaceĩ* from Belize, *S. duffyi* from Jamaica, and *S. rathbunae* from Panama (50). For a meaningful comparison of historical demographics, we included non-eusocial species that showed stable or increasing populations from the same three sites (50).

### Double digest restriction-site associated DNA sequencing

We extracted genomic DNA using several walking legs from alcohol-preserved specimens using Qiagen DNAeasy Tissue Kits. Extracted DNA was quantified using a Qubit 3.0 Fluorometer with the dsDNA HS assay (ThermoFisher Scientific) and visualized on 2% agarose gels. We followed the protocol in Peterson et al. (2012) using the restriction enzymes EcoRI and MspI. We used the recommended size selection criterion (338bp – 414bp) based on the estimated genome sizes of each species, which ranged from approximately 6.49 to 20.74 pg (where 1 pg = 978 Mbp) (79). Briefly, we digested 1,000 ng of genomic DNA with EcoRI and MspI (New England Biolabs) and cleaned the digested DNA using Agencourt AMPure XP beads (Beckman Coulter Life Sciences). We ligated the double-digested DNA with barcoded adaptors that were 5-fold in excess to prevent the formation of chimeras. We pooled and bead-cleaned barcoded samples before size selection using a Pippin Prep and a dye-free 2% Agarose gel cassette with internal standards (CDF2010, Sage Science). We performed a 10-cycle PCR using a Phusion PCR Kit according to the manufacturer’s protocol (New England Biolabs) with multiplexed primers and adjusted PCR products to 10 μM for sequencing on either an Illumina HiSeq2500 (125 bp paired-end, New York Genome Center) or an Illumina HiSeq 3000 (150 bp paired-end, Center for Genome Research and Biocomputing at Oregon State University).

We used *Stacks* v2.1 (80) to demultiplex and clean raw reads (*process_radtags: reads with phred score <10 were discarded*) and perform *de novo* mapping of paired-end reads (*denovo_map.pl* with the parameters T = 6, m = 3, M = 3, and n = 2). We adjusted the parameter *r* (0.5 – 0.875) in *populations* in the *Stacks* pipeline to obtain around 20,000 SNPs per species (19,778 – 52,898).

### Genetic diversity and kinship

We calculated several statistics to summarize the genetic diversity at the individual level, and to quantify the levels of inbreeding and kinship among females, each from different sponges within the same collection site. First, we estimated observed heterozygosity (H_obs_) for each sample from *populations* in the *Stacks* pipeline, which is essentially the number of heterozygous sites divided by the total number of fixed and variant sites for each sample, and took the mean H_obs_ for each species. We also calculated H_obs_ based on variant sites only. We calculated the averaged inbreeding coefficient (F) across individuals for each species using *VCFtools* v0.1.16 (81). F measures the probability that two alleles at a given locus are identical by descent. An individual is fully inbred when F = 1 and fully outbred when F = 0 (82). Finally, we used *King* v2.1.6 (83) to calculate the genome-wide relatedness between each female within a species. We took the mean kinship coefficient between pairs of individuals (one female per sponge) for each species. For each genetic statistic (mean H_obs_, F, and mean kinship coefficient), we performed phylogenetic mixed model regressions using the R package *MCMCglmm* v2.29 (63) to test whether they differed according to the form of social organization (pair-forming vs. communal breeding vs. eusocial) or eusociality (eusocial vs. non-eusocial), while controlling for phylogenetic non-independence between species. We used a Bayesian consensus tree of *Synalpheus* species, constructed with 16S, 18S, and COI sequence data (64). We used weakly informative priors (variance parameters, V = 1, degree of belief, nu = 0.002), but more or less informative priors yielded the same results (nu = 0.02 or 0.0002). We ran 2,000,000 Markov Chain Monte Carlo (MCMC) iterations with 50,000 iterations of burn-in and a thinning interval of 250. Pairwise differences between categories of sociality or eusociality were tested by comparing the difference in posterior distributions of each category (e.g., eusocial – communal breeding) and calculating a pMCMC value as twice the probability that the posterior distribution of the difference is above or below zero (63).

### Demographic inference

For pairs of individuals that were closer than 3^rd^-degree-related (> 25%, based on kinship coefficients > 0.089), we randomly removed one individual using *VCFtools* v0.1.16 (81). We removed two individuals from *S. rathbunae* and one individual each from *S. agelas, S. carpenteri*, and *S. chacei*. We then extracted one random SNP per locus using the script *vcfparser.py* (84) to generate a vcf file for demographic inference analysis using *MOMI2* v2.1.16 (23, 24). For comparative purposes, we chose to infer N_e_ from T fixed time ranges across G generations for all species. Initial N_e_ was set at 1e+6 and maximum N_e_ at 1e+21 for all species. The value of initial N_e_ was based on the mean N_e_ across species from single N_e_ models (See Supporting Information). The value of maximum N_e_ was based on running multiple models with different parameter values and finding the value that yielded the highest log-likelihood across species (See Supporting Information). We ran 300 random starts to find the model with the highest log-likelihood. We further ran 300 bootstraps, each with 50 random starts to find the model with the highest log-likelihood support. Preliminary analyses showed that estimating N_e_ at fixed time ranges (e.g., 1-20, 21-40, 41-60, and 61-80 thousand generations) gave better model support than estimating N_e_ at fixed times (e.g., 20, 40, 60, and 80 thousand generations) (see Supporting Information: Fig. S4).

To find the best specification of model parameters T (number of time ranges) and G (maximum number of generations), we ran seven models that differed in T (4, 6, and 8) and G (60, 80, and 100 thousand). We did not estimate N_e_ changes that were less than 10,000 generations apart because our preliminary analysis showed that the order of magnitude that we can estimate change among the 12 species was about 10,000 generations (See Supporting Information). For each species, we compared the support for each parameter set using delta AIC (62). The model with delta AIC = 0 had the best support and models with delta AIC < 3 were considered to be equally-supported best models. We chose the model with the parameter set that was most frequently supported across species under the criteria of delta AIC < 3 (Table S5).

### Phylogenetic comparative analysis

From the best fit model with the highest log-likelihood, we used the N_e_ values estimated across 10,000 – 100,000 generations for each species. We calculated mean N_e_ across time (mean N_e_), as well as three population stability metrics for comparative analyses: (i) the coefficient of variation of N_e_ based on the log-normal distribution (CV); (ii) the ratio of minimum N_e_ to mean N_e_ (min/mean N_e_); and (iii) the number of times where two consecutive N_e_ estimates had less than one order of magnitude difference (no. of <1-order change). We did not include the N_e_ estimate at time 0 (i.e. the most recent N_e_) in our analysis because this estimate was strongly correlated with the initial N_e_ that we specified in *MOMI2* (data not shown). Further, demographic inference based on ddRAD may not reflect recent population changes (31) unless sample size per population was very large (> 30) (42, 85). We used the Shapiro-Wilk’s test to check for normality, ultimately log-transforming mean N_e_, CV, and min/mean N_e_.

We performed phylogenetic mixed model regressions using *MCMCglmm* (63) as described above. Briefly, we tested whether the four metrics of population size and stability differed according to the form of social organization (pair-forming vs. communal breeding vs. eusocial) or eusociality (eusocial vs. non-eusocial), while controlling for phylogenetic non-independence between species and treating collection site as a random factor. We performed the same analyses using the demographic metrics calculated from the median of 300 bootstraps N_e_ estimates. We also performed a similar analysis using the eusociality index (E = 1 - ((2 × NOF) / CS) where CS, colony size, is the total number of individuals in a sponge and NOF is the number of ovigerous females in a sponge based on data from (64), which captures the degree of reproductive skew within a group and has been used to measure the level of eusociality in *Synalpheus* and other invertebrates (7, 65).

## Acknowledgements

S.T.C.C. was supported by the Simons Foundation via the Life Sciences Research Foundation. S.E.H. was supported by the Frontiers of Science Fellowship at Columbia University. D.R.R. was supported by the US National Science Foundation (IOS-1121435, IOS-1257530, IOS-1439985). This work made use of the high performance computing resources from Columbia University.

## Author Contributions

D.R.R and S.E.H. designed the study. S.T.C.C. and S.E.H. performed the experiment, conducted analyses, and drafted the manuscript. All authors collected specimens and revised the manuscript.

The authors declare no conflict of interest.

## Data Availability

Supporting data is available from the Supporting Information. Sequence reads were deposited in NCBI’s Sequence Read Archive (Accession numbers: SAMN14351547 - SAMN14351641). Additional data is available in Dryad.

