## Supplementary Information for "Demographic inference provides insights into the extirpation and ecological dominance of eusocial snapping shrimps"

**This PDF file includes:**

Supplementary methods

Tables S1 to S6

Figures S1 to S6

### 1. SUPPLEMENTARY METHODS

**Constant population-size model.** The time to coalescence is roughly $N_e$ generations for any population. This value approximates the order of magnitude at which changes in population size can be made. To identify the order of magnitude that $N_e$ change can be estimated from each of the 12 *Synalpheus* species, we ran a constant $N_e$ model for each species using *MOMI2*. The mean $N_e$ from all species was 1,049,317 (range = 28,219 – 4,222,658; median = 770,730), so the minimum allowable change is ~1,000K / 100 = ~10K generations. This means, for example, that estimating eight $N_e$ across 80,000 generations is acceptable, but estimating 16 $N_e$ across 80,000 generations is not.

**Determining the best maximum $N_e$ value.** The maximum $N_e$ value for the demographic models in *MOMI2* was set to be arbitrarily high (1e+21) to avoid constraining estimates of $N_e$. However, we investigated how different values maximum $N_e$ would affect the likelihoods of the models and our conclusions. With a model that estimated $N_e$ four times across 100,000 generations, we used different values of maximum $N_e$ ranging from 1e+4 to 1e+21. For each species, we ranked the models by the log-likelihood and used the model (and the corresponding value of maximum $N_e$) for subsequent analysis.

We found that the model with maximum $N_e$ of 1e+21 was among the best models with delta AIC <1 for each of the 12 species. However, across species, model with maximum $N_e$ of 1e+21 did not always have the highest log-likelihood among models that were equally supported based on AIC. Therefore, we extracted the model with the highest log-likelihood in each species, which had different maximum $N_e$ for each species. From each of these models, we calculated mean $N_e$, coefficient of variation of $N_e$ (CV), min/mean $N_e$, and the no. of <1-order difference in $N_e$ values. We performed phylogenetic mixed model regressions using the four metrics of population size and stability to test whether they differed according to the form of social organization (pair-forming vs. communal breeding vs. eusocial). We also calculated the demographic metrics for each of the 300 bootstrap results and ran the regression analyses with the median of the demographic metrics calculated from the bootstrap results.

We found that the relationships between social organization and mean $N_e$, CV, and min/mean $N_e$ remained the same despite not using models for maximum $N_e$ of 1e+21 for all species (Figure S**). Results from the median of bootstrap metrics also showed a consistent pattern (Figure S**). Therefore, regardless of the parameter used for the demographic models, we observed the same pattern: eusocial species had lower and more stable $N_e$ through time than non-eusocial species.

### 2. SUPPLEMENTARY TABLES

**Table S1**. Metadata for each *Synalpheus* species. Minimum percent of population per locus shows the percent of sampled individuals required to be genotyped at any particular ddRAD tag locus. Number of random SNPs shows the final number of SNPs used in the analysis when one random SNP per ddRAD tag locus was kept.

| ***Synalpheus* species** | **Social organization** | **Collection country** | **Collection years** | **Number of samples** | **Number of kin samples removed** | **Minimum percent of population per locus** | **Median number of reads per sample** | **Number of SNPs from Stacks** | **Number of random SNPs** |
|---|---|---|---|---|---|---|---|---|---|
| *agelas* | Pair | Jamaica | 2008 | 8 | 1 | 0.5 | 1,209,211 | 23,276 | 14,118 |
| *bousfieldi* | Pair | Belize | 2005, 2009 | 8 | | 0.63 | 3,126,365 | 23,246 | 12,618 |
| *brooksi* | Eusocial | Belize | 2005, 2009 | 8 | | 0.5 | 1,630,730 | 22,520 | 26,148 |
| *carpenteri* | Communal | Jamaica | 2008 | 8 | 1 | 0.5 | 2,326,561 | 19,778 | 12,816 |
| *chacei* | Eusocial | Belize | 2004 | 9 | 1 | 0.56 | 2,554,968 | 29,894 | 26,114 |
| *dardeaui* | Communal | Belize | 2005, 2009 | 8 | | 0.5 | 812,256 | 42,480 | 21,143 |
| *duffyi* | Eusocial | Jamaica | 2008, 2012 | 7 | | 0.58 | 4,503,672 | 35,329 | 13,065 |
| *herricki* | Communal | Barbados | 2008 | 8 | | 0.5 | 1,944,451 | 21,593 | 23,221 |
| *idios* | Communal | Barbados | 2008 | 8 | | 0.75 | 3,960,982 | 25,337 | 18,558 |
| *longicarpus small* | Pair | Panama | 2007, 2008, 2009 | 8 | | 0.5 | 4,316,153 | 52,898 | 24,707 |
| *rathbunae* | Eusocial | Panama | 2007, 2008 | 8 | 2 | 0.88 | 6,086,252 | 39,548 | 11,890 |
| *yano* | Communal | Panama | 2007, 2008, 2009 | 8 | | 0.75 | 4,950,380 | 22,300 | 13,702 |

**Supplementary Table S2.** Metadata for each *Synalpheus* sample.

| ***Synalpheus* species** | **Sample names** | **Number of paired-end reads** | **Collection country** | **Collection year** | **SRA accession number** |
|---|---|---|---|---|---|
| *agelas* | JAM2008-014-002_agelas | 1798247 | Jamaica | 2008 | SAMN14351547 |
| *agelas* | JAM2008-020-001_agelas | 110557 | Jamaica | 2008 | SAMN14351548 |
| *agelas* | JAM2008-020-003_agelas | 211880 | Jamaica | 2008 | SAMN14351549 |
| *agelas* | JAM2008-030-001_agelas | 4844719 | Jamaica | 2008 | SAMN14351550 |
| *agelas* | JAM2008-044-001_agelas | 3018677 | Jamaica | 2008 | SAMN14351551 |
| *agelas* | JAM2008-056-001_agelas | 6349613 | Jamaica | 2008 | SAMN14351552 |
| *agelas* | JAM2008-061-006_agelas | 301232 | Jamaica | 2008 | SAMN14351553 |
| *agelas* | JAM2008-085-003_agelas | 620174 | Jamaica | 2008 | SAMN14351554 |
| *bousfieldi* | CBC2005_037_006_002 | 789950 | Belize | 2005 | SAMN14351555 |
| *bousfieldi* | CBC2005_037_006_003 | 219946 | Belize | 2005 | SAMN14351556 |
| *bousfieldi* | CBC2009_024_001 | 8009135 | Belize | 2009 | SAMN14351557 |
| *bousfieldi* | CBC2009_024_005 | 3648579 | Belize | 2009 | SAMN14351558 |
| *bousfieldi* | CBC2009_036_003 | 5340429 | Belize | 2009 | SAMN14351559 |
| *bousfieldi* | CBC2009_036_006 | 3965235 | Belize | 2009 | SAMN14351560 |
| *bousfieldi* | CBC2009_066_003 | 22093 | Belize | 2009 | SAMN14351561 |
| *bousfieldi* | CBC2009_088_002 | 2604151 | Belize | 2009 | SAMN14351562 |
| *brooksi* | CBC2005_002_008 | 893878 | Belize | 2005 | SAMN14351563 |
| *brooksi* | CBC2005_014_002 | 1360106 | Belize | 2005 | SAMN14351564 |
| *brooksi* | CBC2005_031_006_001 | 1901353 | Belize | 2005 | SAMN14351565 |
| *brooksi* | CBC2005_031_006_002 | 933739 | Belize | 2005 | SAMN14351566 |
| *brooksi* | CBC2005_032_001 | 240944 | Belize | 2005 | SAMN14351567 |
| *brooksi* | CBC2009_008_004 | 4956979 | Belize | 2009 | SAMN14351568 |
| *brooksi* | CBC2009_040_002 | 5855667 | Belize | 2009 | SAMN14351569 |
| *brooksi* | CBC2009_060_001 | 2283699 | Belize | 2009 | SAMN14351570 |
| *carpenteri* | JAM2008-010-001_carpenteri | 2351111 | Jamaica | 2008 | SAMN14351571 |
| *carpenteri* | JAM2008-013-001_carpenteri | 61660 | Jamaica | 2008 | SAMN14351572 |
| *carpenteri* | JAM2008-020-006_carpenteri | 2302011 | Jamaica | 2008 | SAMN14351573 |
| *carpenteri* | JAM2008-026-001_carpenteri | 3144025 | Jamaica | 2008 | SAMN14351574 |
| *carpenteri* | JAM2008-030-004_carpenteri | 3040950 | Jamaica | 2008 | SAMN14351575 |
| *carpenteri* | JAM2008-035-001_carpenteri | 3679030 | Jamaica | 2008 | SAMN14351576 |
| *carpenteri* | JAM2008-038-001_carpenteri | 1048414 | Jamaica | 2008 | SAMN14351577 |
| *carpenteri* | JAM2008-039-001_carpenteri | 85799 | Jamaica | 2008 | SAMN14351578 |
| *chacei* | CBC2004-005-003_chacei | 5254142 | Belize | 2004 | SAMN14351579 |
| *chacei* | CBC2004-012-001_chacei | 15358 | Belize | 2004 | SAMN14351580 |
| *chacei* | CBC2004-042-004_chacei | 1979216 | Belize | 2004 | SAMN14351581 |
| *chacei* | CBC2004-044-001_chacei | 2464917 | Belize | 2004 | SAMN14351582 |

| | | | | | |
|---|---|---|---|---|---|
| *chacei* | CBC2004-056-001_chacei | 2645019 | Belize | 2004 | SAMN14351583 |
| *chacei* | CBC2004-060-002_chacei | 4068533 | Belize | 2004 | SAMN14351584 |
| *chacei* | CBC2004-061-002_chacei | 98190 | Belize | 2004 | SAMN14351585 |
| *chacei* | CBC2004-062-001_chacei | 5001232 | Belize | 2004 | SAMN14351586 |
| *dardeaui* | CBC2005_002_014 | 457363 | Belize | 2005 | SAMN14351587 |
| *dardeaui* | CBC2005_014_017 | 99755 | Belize | 2005 | SAMN14351588 |
| *dardeaui* | CBC2005_030_003 | 1167149 | Belize | 2005 | SAMN14351589 |
| *dardeaui* | CBC2005_033_001 | 326744 | Belize | 2005 | SAMN14351590 |
| *dardeaui* | CBC2005_042_004 | 27545 | Belize | 2005 | SAMN14351591 |
| *dardeaui* | CBC2009_036_009 | 1797615 | Belize | 2009 | SAMN14351592 |
| *dardeaui* | CBC2009_039_001 | 4192164 | Belize | 2009 | SAMN14351593 |
| *dardeaui* | CBC2009_040_008 | 2727844 | Belize | 2009 | SAMN14351594 |
| *duffyi* | JAM2008-009-001_duffyi | 1754737 | Jamaica | 2008 | SAMN14351595 |
| *duffyi* | JAM2008-012-004_duffyi | 772383 | Jamaica | 2008 | SAMN14351596 |
| *duffyi* | JAM2008-050-001_duffyi | 4814358 | Jamaica | 2008 | SAMN14351597 |
| *duffyi* | JAM2008-075-008_duffyi | 2965308 | Jamaica | 2008 | SAMN14351598 |
| *duffyi* | JAM2012-135-003_duffyi | 1944451 | Jamaica | 2012 | SAMN14351599 |
| *duffyi* | JAM2012-165-001_duffyi | 2470843 | Jamaica | 2012 | SAMN14351600 |
| *duffyi* | JAM12_12307_duffyi | 1808483 | Jamaica | 2012 | SAMN14351601 |
| *herricki* | BR08_1501_herricki | 778758 | Barbados | 2008 | SAMN14351602 |
| *herricki* | BR08_1502_01_herricki | 1071256 | Barbados | 2008 | SAMN14351603 |
| *herricki* | BR08_1502_2A_herricki | 47679 | Barbados | 2008 | SAMN14351604 |
| *herricki* | BR08_8101_herricki | 450715 | Barbados | 2008 | SAMN14351605 |
| *herricki* | BR08_8102_herricki | 2793247 | Barbados | 2008 | SAMN14351606 |
| *herricki* | BR08_8103_herricki | 1470594 | Barbados | 2008 | SAMN14351607 |
| *herricki* | BR08_8104_herricki | 1543520 | Barbados | 2008 | SAMN14351608 |
| *herricki* | BR08_8105_01_herricki | 652489 | Barbados | 2008 | SAMN14351609 |
| *idios* | BR08_101_01_idios | 5030924 | Barbados | 2008 | SAMN14351610 |
| *idios* | BR08_101_03_idios | 368600 | Barbados | 2008 | SAMN14351611 |
| *idios* | BR08_101_04_idios | 2107983 | Barbados | 2008 | SAMN14351612 |
| *idios* | BR08_104_02_idios | 1995601 | Barbados | 2008 | SAMN14351613 |
| *idios* | BR08_105_02_idios | 2891040 | Barbados | 2008 | SAMN14351614 |
| *idios* | BR08_105_03_idios | 5755017 | Barbados | 2008 | SAMN14351615 |
| *idios* | BR08_105_04_idios | 6666596 | Barbados | 2008 | SAMN14351616 |
| *idios* | BR08_105_05_idios | 5667473 | Barbados | 2008 | SAMN14351617 |
| *longicarpus small* | P2007_012_001 | 8466716 | Panama | 2007 | SAMN14351618 |
| *longicarpus small* | P2007_032_003 | 2398138 | Panama | 2007 | SAMN14351619 |
| *longicarpus small* | P2007_053_001 | 4337645 | Panama | 2007 | SAMN14351620 |
| *longicarpus small* | P2007_071_001 | 4294660 | Panama | 2007 | SAMN14351621 |

| | | | | | |
|---|---|---|---|---|---|
| *longicarpus small* | P2008_080_001 | 6598208 | Panama | 2008 | SAMN14351622 |
| *longicarpus small* | P2008_134_002 | 3239101 | Panama | 2008 | SAMN14351623 |
| *longicarpus small* | P2009_039_002 | 12309 | Panama | 2009 | SAMN14351624 |
| *longicarpus small* | P2009_092_002 | 4505769 | Panama | 2009 | SAMN14351625 |
| *rathbunae* | P08_129_03_1 | 9229654 | Panama | 2008 | SAMN14351626 |
| *rathbunae* | P2007_039_001_018 | 1273562 | Panama | 2007 | SAMN14351627 |
| *rathbunae* | P2007_039_002_002 | 11970422 | Panama | 2007 | SAMN14351628 |
| *rathbunae* | P2008_115_006 | 6513776 | Panama | 2008 | SAMN14351629 |
| *rathbunae* | P2008_117_003 | 443487 | Panama | 2008 | SAMN14351630 |
| *rathbunae* | P2008_129_003_002 | 5658728 | Panama | 2008 | SAMN14351631 |
| *rathbunae* | P2008_129_006_001 | 10400647 | Panama | 2008 | SAMN14351632 |
| *rathbunae* | P2008_131_004 | 5552645 | Panama | 2008 | SAMN14351633 |
| *yano* | BDT2011_209_004_002 | 4570682 | Panama | 2012 | SAMN14351634 |
| *yano* | P2007_013_003 | 15635814 | Panama | 2007 | SAMN14351635 |
| *yano* | P2007_035_003 | 5980780 | Panama | 2007 | SAMN14351636 |
| *yano* | P2007_060_003 | 5330077 | Panama | 2007 | SAMN14351637 |
| *yano* | P2008_003_011 | 5401187 | Panama | 2008 | SAMN14351638 |
| *yano* | P2008_017_002 | 3090352 | Panama | 2008 | SAMN14351639 |
| *yano* | P2008_090_011 | 3881328 | Panama | 2008 | SAMN14351640 |
| *yano* | P2009_020_001 | 618386 | Panama | 2009 | SAMN14351641 |

**Table S3.** Population genetic statistics for each *Synalpheus* species. Mean individual observed heterozygosity ($H_{obs}$) is the percentage of heterozygous sites across all sites or variant site only. Pair: pair-living, communal: communal breeding. Kinship coefficients of 0.044 to 0.088 indicates 3rd degree relatedness, coefficients of 0.088 to 0.18 indicates 2nd degree relatedness, coefficients of 0.18 to 0.35 indicates 1st degree relatedness, coefficients > 0.35 indicates duplicate/MZtwins, and negative values indicate unrelated relationships.

| ***Synalpheus* species** | **Social organization** | **$H_{obs}$ (all sites)** | **$H_{obs}$ (variant sites)** | **Inbreeding coefficient ($F_{IS}$)** | **Mean kinship coefficient** |
|---|---|---|---|---|---|
| *agelas* | Pair | 0.22 | 0.44 | 0.29 | -2.05 |
| *bousfieldi* | Pair | 0.20 | 0.50 | 0.32 | -2.74 |
| *longicarpus small* | Pair | 0.22 | 0.60 | 0.12 | -1.70 |
| *carpenteri* | Communal | 0.23 | 0.45 | 0.30 | -1.46 |
| *dardeaui* | Communal | 0.23 | 0.34 | 0.26 | -0.53 |
| *herricki* | Communal | 0.22 | 0.67 | 0.14 | -0.90 |
| *idios* | Communal | 0.18 | 0.19 | 0.54 | -0.71 |
| *yano* | Communal | 0.20 | 0.28 | 0.45 | -0.93 |
| *brooksi* | Eusocial | 0.19 | 0.38 | 0.13 | -0.14 |
| *chacei* | Eusocial | 0.20 | 0.31 | 0.16 | -0.22 |
| *duffyi* | Eusocial | 0.22 | 0.45 | 0.21 | -1.19 |
| *rathbunae* | Eusocial | 0.26 | 0.45 | 0.14 | -0.26 |

**Table S4.** Parameters estimated from phylogenetic mixed models which tested the effect of sociality (Pair: pair-living, Communal: communal breeding, and Eusocial: eusociality) on genetic diversity and kinship coefficients between 12 species of *Synalpheus* species. Asterisks (*) indicate pMCMC < 0.05 in pairwise comparison between forms of social organization and between eusocial and non-eusocial (pair-forming and communal breeding) species.

| Predictor variable | Response variable | delta DIC | Comparison | Posterior mean | Highest probability density | Effective sample size | pMCMC | |
|---|---|---|---|---|---|---|---|---|
| Sociality | $H_{obs}$ (all sites) | -4.960 | Eusocial - Communal | -0.00021 | -0.042 | 0.041 | 0.987 | |
| | | | Eusocial - Pair | -0.00044 | -0.046 | 0.041 | 0.985 | |
| | | | Communal - Pair | -0.00023 | -0.044 | 0.040 | 0.996 | |
| | | | Eusocial - Non-eusocial | -0.00033 | -0.039 | 0.036 | 0.979 | |
| Sociality | $H_{obs}$ (variant sites) | 3.110 | Eusocial - Communal | 0.015 | -0.207 | 0.229 | 0.885 | |
| | | | Eusocial - Pair | -0.130 | -0.377 | 0.099 | 0.245 | |
| | | | Communal - Pair | -0.145 | -0.386 | 0.081 | 0.186 | |
| | | | Eusocial - Non-eusocial | -0.057 | -0.265 | 0.130 | 0.511 | |
| Sociality | $F_{IS}$ | 8.800 | Eusocial - Communal | -0.349 | -0.647 | -0.022 | 0.032 | * |
| | | | Eusocial - Pair | -0.186 | -0.511 | 0.140 | 0.241 | |
| | | | Communal - Pair | 0.162 | -0.154 | 0.472 | 0.266 | |
| | | | Eusocial - Non-eusocial | -0.267 | -0.546 | 0.019 | 0.062 | |
| Sociality | Kinship coefficient | 32.080 | Eusocial - Communal | 0.702 | -0.007 | 1.526 | 0.069 | |
| | | | Eusocial - Pair | 1.978 | 1.177 | 2.781 | 0.0003 | * |
| | | | Communal - Pair | 1.276 | 0.609 | 1.988 | 0.004 | * |
| | | | Eusocial - Non-eusocial | 1.340 | 0.625 | 2.036 | 0.002 | * |

**Table S5**. Count of species that supported a model based on delta AIC ≤ 2 to ≤ 3. The models differed by the number of $N_e$ estimates (T) and the maximum number of generations (in thousand) (G). The model that had the best support across species is highlighted in bold.

| | | **Count of species supporting model with** | |
|---|---|---|---|
| **Number of $N_e$ estimates (T)** | **Maximum number of generations (G)** | **delta AIC ≤ 2** | **delta AIC ≤ 3** |
| 4 | 60 | 6 | 7 |
| 4 | 80 | 7 | 7 |
| **4** | **100** | **8** | **8** |
| 6 | 60 | 2 | 2 |
| 6 | 80 | 2 | 2 |
| 6 | 100 | 0 | 0 |
| 8 | 80 | 2 | 2 |
| 8 | 100 | 0 | 0 |

**Table S6.** Parameters estimated from phylogenetic mixed models that tested the effect of social organization (Pair: pair-living, Communal: communal breeding, and Eusocial: eusociality) on metrics of population size and stability for 12 species of *Synalpheus* species. CV: coefficient of variation of $N_e$.

| Predictor | Response | delta DIC | Comparison | Posterior mean | 95% probability density | Effective sample size | pMCMC |
|---|---|---|---|---|---|---|---|
| Sociality | mean $N_e$ | 35.42 | Eusocial - Communal | -8.24 | -16.26 – 7.96 | 8621 | 0.038 |
| | mean $N_e$ | | Eusocial - Pair | -8.54 | -16.97 – 9.96 | 7047 | 0.078 |
| | mean $N_e$ | | Communal - Pair | -0.31 | -8.86 – 8.48 | 7249 | 0.92 |
| | CV | 16.14 | Eusocial - Communal | -44.48 | -94.91 – 4.46 | 7800 | 0.071 |
| | CV | | Eusocial - Pair | -32.91 | -89.11 – 21.39 | 7427 | 0.21 |
| | CV | | Communal - Pair | 11.57 | -41.32 – 65.48 | 6832 | 0.66 |
| | min/mean $N_e$ | 25.96 | Eusocial - Communal | 7.30 | -1.497 – 16.41 | 7832 | 0.095 |
| | min/mean $N_e$ | | Eusocial - Pair | 6.92 | -2.92 – 16.42 | 6557 | 0.15 |
| | min/mean $N_e$ | | Communal - Pair | -0.38 | -10.63 – 9.23 | 7016 | 0.96 |
| | no. <1-order change | 13.30 | Eusocial - Communal | 0.887 | 0.008 – 1.86 | 7659 | 0.056 |
| | no. <1-order change | | Eusocial - Pair | 0.525 | -0.40 – 1.45 | 7800 | 0.23 |
| | no. <1-order change | | Communal - Pair | -0.362 | -1.28 – 0.52 | 7800 | 0.38 |
| Eusociality | mean $N_e$ | 31.92 | Eusocial - Non-eusocial | 8.614 | 1.82 – 15.54 | 8672 | 0.021 |
| | CV | 29.62 | Eusocial - Non-eusocial | 40.908 | -0.66 – 83.05 | 8279 | 0.055 |
| | min/mean $N_e$ | 26.54 | Eusocial - Non-eusocial | -7.427 | -15.01 – 0.097 | 7507 | 0.058 |
| | no. <1-order change | 14.11 | Eusocial - Non-eusocial | -0.698 | -1.45 – 0.067 | 8270 | 0.076 |
| Eusociality index | mean $N_e$ | 30.83 | | -7.11 | -12.65 to -1.33 | 7268 | 0.025 |
| | CV | 67.73 | | -35.85 | -66.65 to -5.25 | 8135 | 0.031 |
| | min/mean $N_e$ | 58.02 | | 7.22 | 2.12 to 12.53 | 7800 | 0.015 |
| | no. <1-order change | -0.46 | | 0.17 | -0.62 to 8.94 | 7800 | 0.63 |

Additional data is available in Dryad: Model output of $N_e$ and time of the best model and the bootstrap's median and 95% confidence interval.

### 3. SUPPLEMENTARY FIGURES

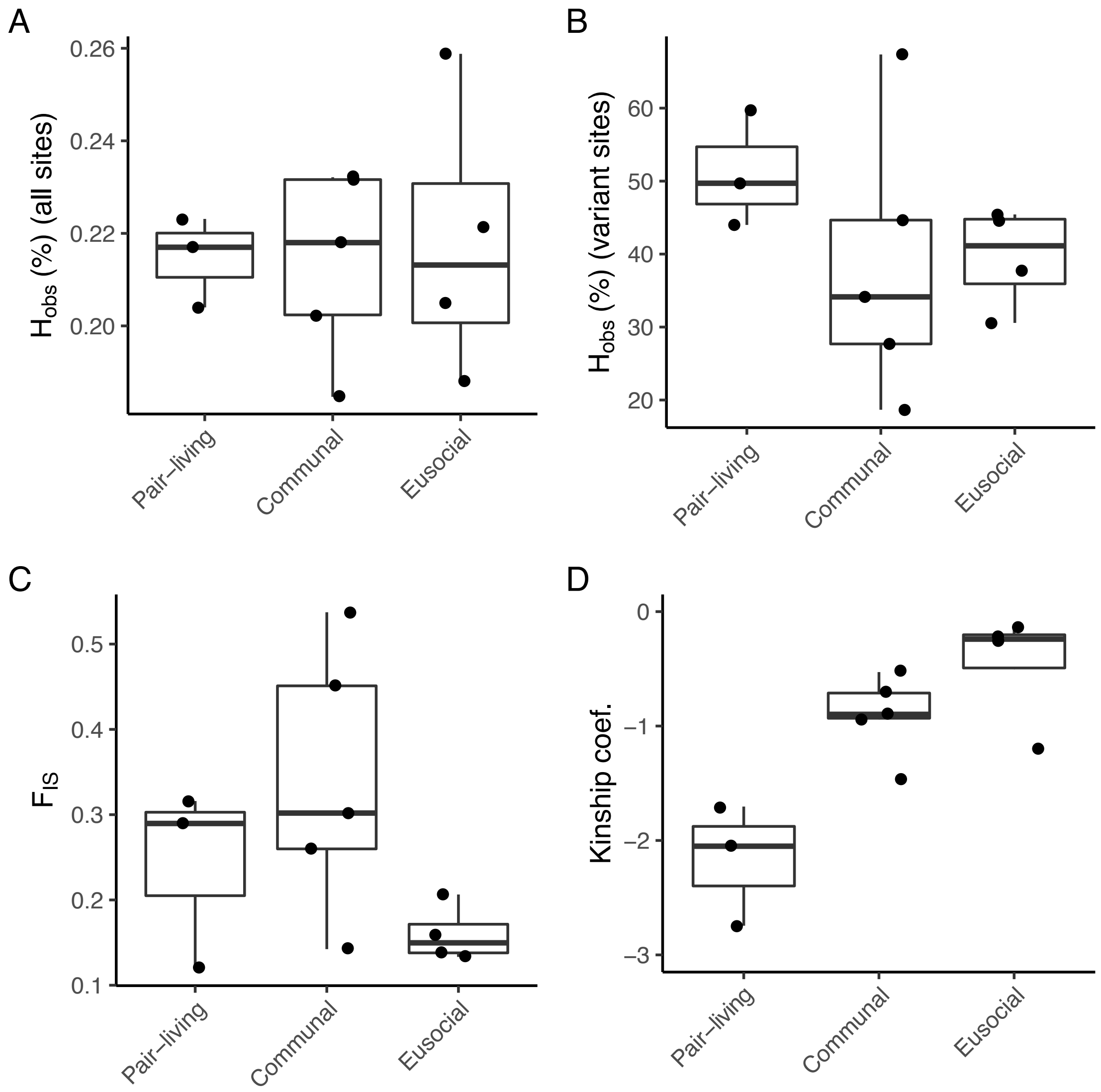


**Fig. S1**. Genetic diversity and kinship of *Synalpheus* species exhibiting different forms of social organization. (A) Mean individual observed heterozygosity ($H_{obs}$) based on all sites, which is the number of heterozygous sites divided by the total number of fixed and variant sites for each sample. (B) $H_{obs}$ based on variant site. (C) Averaged inbreeding coefficients ($F_{IS}$) across samples. A $F_{IS}$ of 1 indicates fully inbred, whereas -1 indicates fully outbred. (D) Mean kinship coefficient between individuals from different sponges in the same collection site. Kinship coefficients of 0.044 to 0.088 indicates 3rd degree relatedness, coefficients of 0.088 to 0.18 indicates 2nd degree relatedness, coefficients of 0.18 to 0.35 indicates 1st degree relatedness, coefficients > 0.35 indicates duplicate/MZtwins, and negative values indicate unrelated relationships.

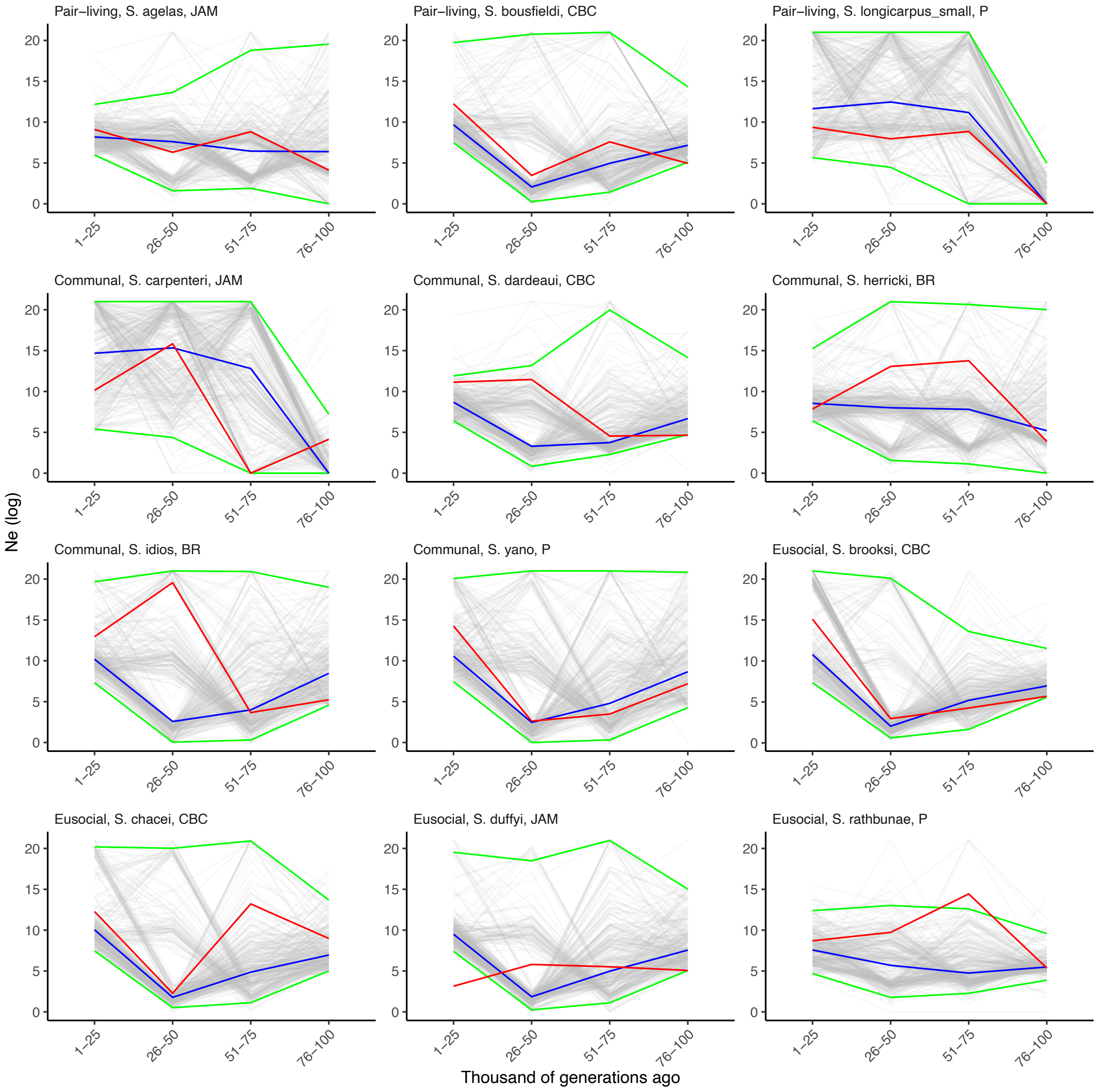


**Fig. S2**. $N_e$ estimates based on the best model (red lines) and 300 bootstraps in gray (blue lines: median of bootstraps, green lines: 95% confidence intervals) in three eusocial, five communal breeding and three pair-living *Synalpheus* species. Letters next to species names indicate locations (BR: Barbados; CBC: Carrie Bow Cay, Belize, J: Jamaica, and P: Panama).

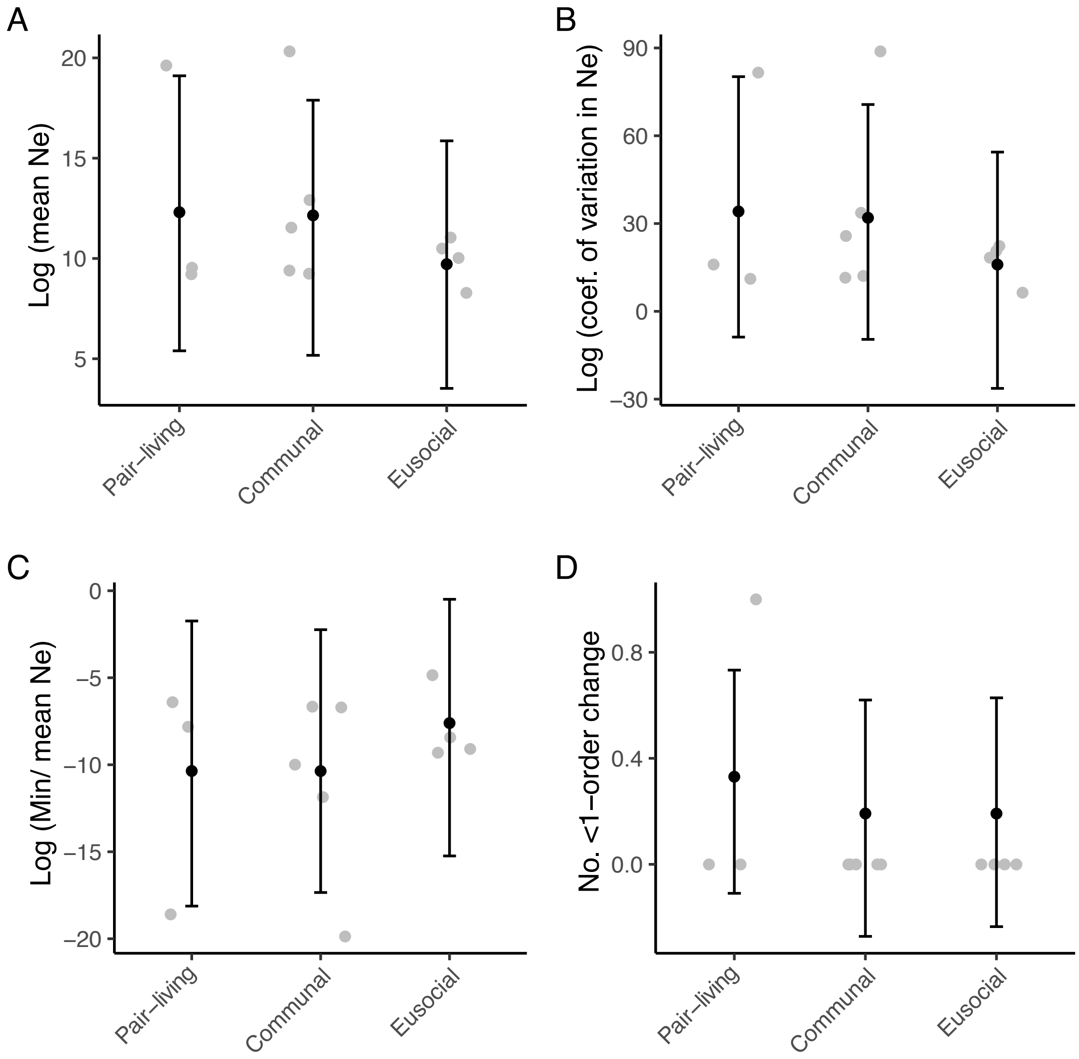


**Fig. S3**. Metrics of population size and stability across 12 *Synalpheus* species exhibiting different forms of social organization based on the median of 300 bootstraps $N_e$ estimates. Grey dots are raw values, black dots are posterior means predicted using Bayesian phylogenetic mixed models, bars indicate the 95% posterior distributions.

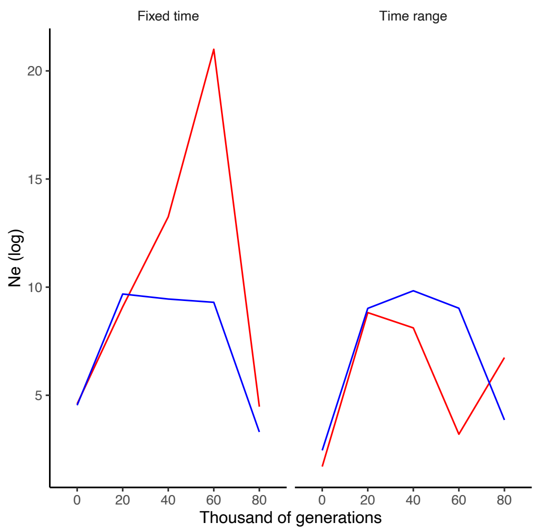


**Fig. S4.** $N_e$ estimates from the best model (red) and median $N_e$ of 100 bootstrap estimates (blue) in a representative species (*S. agelas*) using models with fixed time and the time range. Other species (not shown) showed similar trend in which the best model and bootstrap estimates were more similar in models using the time range than those using fixed time.

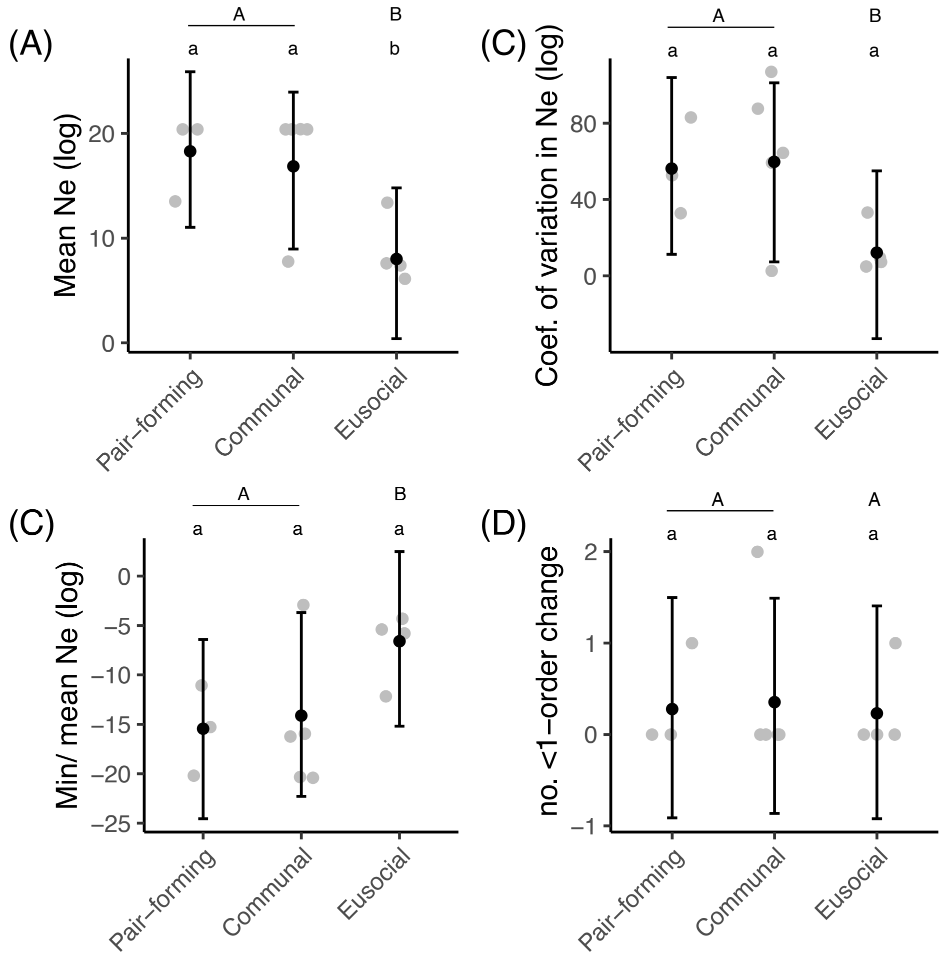


**Fig. S5.** Metrics of population size and stability for 12 *Synalpheus* shrimps exhibiting different forms of social organization. The data is based on model with the highest log-likelihood based on different parameter of maximum $N_e$ in the demographic model. Consistent with the main result from models with maximum $N_e$ of 1e+21, eusocial species have lower mean $N_e$ across time (A), but more stable values of $N_e$ across generation time as indicated by lower CV (B), and higher min/mean $N_e$ (C). The no. <1-order change (D) did not differ between social organization, but was higher in eusocial species based on the bootstrap median results (Figure S6). Grey dots are raw values, black dots are posterior mean predicted using Bayesian phylogenetic mixed model, bars are the 95% posterior distributions. Asterisks (*) indicate pMCMC < 0.05 in pairwise comparison between forms of social organization and between eusocial and non-eusocial (pair-forming and communal breeding) species. pMCMC values are twice the probability that the posterior distribution of the difference is above or below zero.

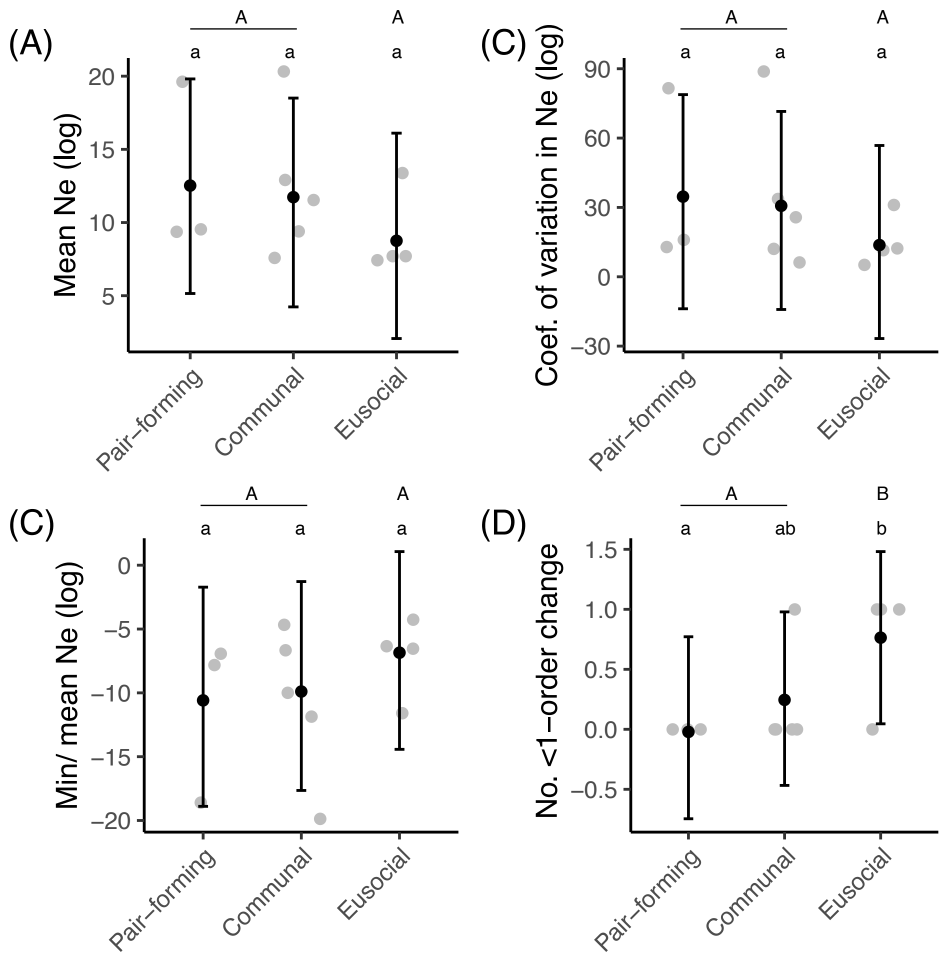


**Fig. S6.** Metrics of population size and stability for 12 *Synalpheus* shrimps exhibiting different forms of social organization. The data is based on model with the highest log-likelihood based on different parameter of maximum $N_e$ in the demographic model and the median of the demographic metrics from each of the 300 bootstrap results. Consistent with the main result from models with maximum $N_e$ if 1e+21, eusocial species have lower mean $N_e$ (A), but more stable values of $N_e$ across generation time as indicated by lower CV (B), higher min/mean $N_e$ (C), and higher no. <1-order change (D). Grey dots are raw values, black dots are posterior mean predicted using Bayesian phylogenetic mixed model, bars are the 95% posterior distributions. Asterisks (*) indicate pMCMC < 0.05 in pairwise comparison between forms of social organization and between eusocial and non-eusocial (pair-forming and communal breeding) species. pMCMC values are twice the probability that the posterior distribution of the difference is above or below zero.
